# Soil microbes mediate the effects of environmental variability on plant invasion

**DOI:** 10.1101/2021.11.01.466853

**Authors:** Xue Zhang, Mark van Kleunen, Chunling Chang, Yanjie Liu

**Affiliations:** Key Laboratory of Wetland Ecology and Environment, Northeast Institute of geography and groecology, Chinese Academy of Sciences, Changchun, 130102, China; University of Chinese Academy of Sciences, Beijing, 100049, China; Ecology, Department of Biology, University of Konstanz, 78464 Konstanz, Germany; Zhejiang Provincial Key Laboratory of Plant Evolutionary Ecology and Conservation, Taizhou University, Taizhou 318000, China

**Keywords:** global change, niche, plant invasion, resource competition, soil biota, trophic level

## Abstract

Many studies indicate that increases in resource variability promote plant invasion. However, it remains unknown to what extent these effects might indirectly be mediated by other organisms. To test this, we grew eight alien species in pot-mesocosms with five different native communities under eight combinations of two nutrient-availability, two nutrient-fluctuation and two soil-microbe treatments. We found that when plants grew in sterilized soil, nutrient fluctuation promoted the dominance of alien plants under low nutrient availability, whereas its effect was minimal under high nutrient availability. However, the opposite pattern was found when plants grew in living soil. Analysis of the soil microbial community suggests that this might reflect that nutrient fluctuation strongly increased the soil fungal pathogen diversity under high nutrient availability, but slightly decreased it under low nutrient availability. Our findings indicate that besides its direct influence, environmental variability could also indirectly affect plant invasion via changes in soil-microbial communities.

## Introduction

Eutrophication, increases in environmental variability and rapid accumulation of alien species are three major characteristics of the Anthropocene ^1–3^. Plant invasions often lead to highly undesirable changes in ecosystem structure and function ^4,5^, and are frequently driven by changes in resource availability and environmental variability. The fluctuating resource hypothesis predicts that a plant community becomes more susceptible to invasion whenever there is a fluctuation (i.e. a temporal increase) in resource availability ^6^. Although it has become a widely accepted theory in ecology, empirical studies that tested it ^7–12^ have found mixed results. For example, whereas Parepa et al. ^7^ found that a nutrient pulse increased the dominance of an invasive plant, Liu et al. ^9^ showed that a pulsed nutrient supply decreased the dominance of invasive alien plants. Therefore, it has become an urgent quest to understand when and how fluctuating resources affect alien plant invasion in native communities.

The fluctuating resources hypothesis rests on the assumption that the native community is resource limited ^6^. Consequently, a resource pulse could lead to shifts in the intensity of alien-native competition, increased dominance of alien species and potential competitive exclusion of the natives over time ^8,13^. When, however, resources are not limiting, fluctuations in resource availability should not benefit the aliens, even if they can respond quicker to resource fluctuations ^8^ or have higher maximum growth rates than natives ^7^. If this is true, it logically follows that if the native community grows under more nutrient-limiting conditions, the positive effect of fluctuating resources on alien plant invasion should be stronger. However, very few studies have tested whether this expectation holds, as most previous studies tested the fluctuating resource hypothesis only under relative high nutrient conditions. This might explain why previous studies found mixed results.

While the fluctuating resource hypothesis explains how environmental variability in general should affect alien plant invasion, it considers only the effect of resource availability. Invasion success of alien plants can also be driven by biotic factors. For example, the enemy release hypothesis predicts that the absence or reduced numbers of enemies, such as microbial pathogens, in their introduced range drives invasion success of alien plants ^14,15^. On the other hand, the missed mutualisms hypothesis predicts that alien plants suffer from the absence or reduced numbers of suitable mutualistic partners, such as arbuscular mycorrhiza fungus (AMF; ^16^. Furthermore, the environmental context (e.g., resource availability) may alter the relative contribution of pathogens versus mutualists to plant performance, and thus shape plant-species coexistence ^17^. Following this logic, we expect that fluctuations in resource availability should not only directly but also indirectly—through changes in soil-microbial communities— influence invasion success of alien plants into native communities.

To test the interactive effects among nutrient availability, resource fluctuations and soil microbes, we did a multispecies experiment (Fig. 1). We grew eight naturalized alien plants as target species in pot-mesocosms with five different synthetic native communities under eight combinations of two nutrient-availability (low *vs* high), two nutrient-fluctuation (constant *vs* pulsed) and two soil-microbe (living *vs* sterilized) treatments. We compared the absolute aboveground biomass production of the alien target species as well as their biomass production relative to the aboveground biomass production of the native competitors. Moreover, we used amplicon DNA sequencing of the soil in the mesocosms of the living soil-microbe treatment to assess how the nutrient treatments affect the soil bacterial and fungal communities. We addressed the following questions: (1) Does the effect of nutrient fluctuations on absolute and relative biomass of alien plants depend on the average nutrient availability level? (2) Does the presence of soil microbes change the effects of increases in nutrient availability and fluctuations therein on the absolute and relative biomass of alien plants?

**Figure 1.**
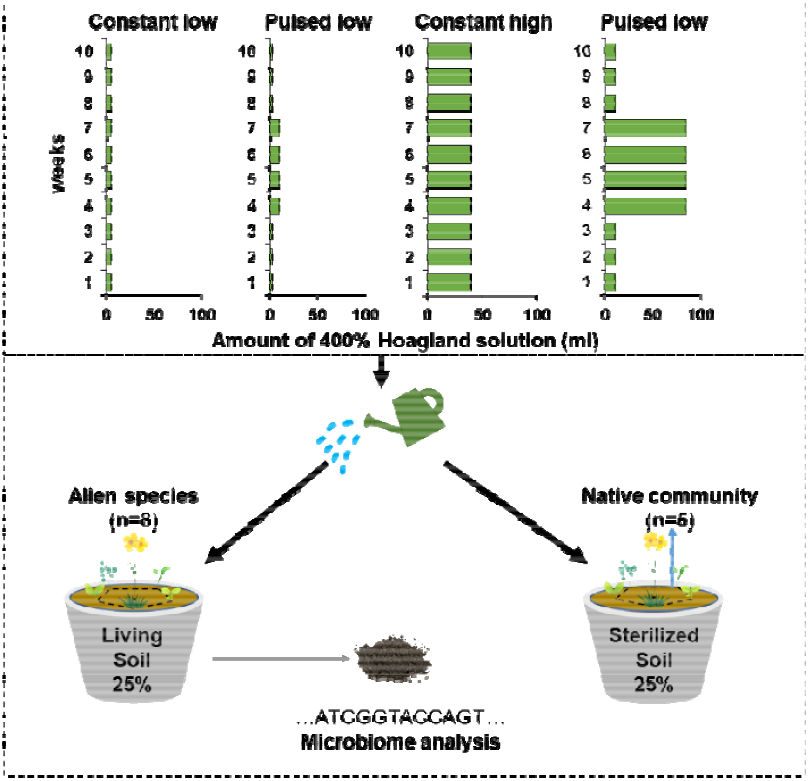
Graphical illustration of the experimental design. The upper panel shows the amounts of nutrient solution supplied to each combination of the nutrient-availability and nutrient-fluctuation treatments during the ten weeks of the experiment. The lower panel shows an overview of the soil-microbe treatment and the locations of the alien target species (plant symbol with yellow flowers) and native community plants in the pots. At the end of the experiment, soil was collected from the pots that had been inoculated with living soil for microbiome analysis.

## Material and Methods

### Study location and species

We conducted our multi-species experiment in a greenhouse of the Northeast Institute of Geography and Agroecology, Chinese Academy of Sciences (43°59’49”N, 125°24’30”E). We selected eight naturalized alien species as targets and 25 native species as native-community members (Table S1). All species are herbaceous plants, and commonly co-occur in many grasslands of China. To cover a wide taxonomic breadth, and thus to increase the generalizability of our findings, we selected the eight alien target species from six genera of three families, and the 25 native species from 22 genera of eleven families (Table S1). We classified the species as naturalized alien or native to China based on the book “The Checklist of the Alien Invasive Plants in China” ^18^ and the Flora of China database (www.efloras.org). Seeds of the study species were collected from wild populations or obtained from commercial seeds companies (Table S1).

### Experimental set-up

From 26 June to 14 August 2020 (Table S1), we sowed the study species separately into plastic trays (19.5 cm × 14.6 cm × 6.5 cm) filled with peat moss as subtrate (Pindstrup Plus, Pindstrup Mosebrug A/S, Denmark). Because the time required for germination were known to differ among the species, we sowed them on different dates (Table S1) to ensure that all species were in similar developmental stages at the start of the experiment. We placed all trays with seeds in a greenhouse under natural light conditions, with a temperature between 20 and 28 °C.

On 30 August 2020, we transplanted similar-sized seedlings of each species into 2.5 L circular plastic pots (top diameter × bottom diameter × height: 18.5 cm × 12.5 cm × 15 cm). We transplanted one seedling of an alien target species in the center of each pot. For each of the eight target species, we transplanted at total of 40 seedlings into 40 pots (i.e., one seedling per pot). We then randomly assigned the 40 pots of each alien target species to be planted with five different native grassland communities (i.e., eight pots per alien target species per native community). We used five native communities to increase generalizability of the results. The communities were created by randomly assigning the 25 native species into five groups of five species (Table S1). We planted one seedling of each native-community member so that each pot included five individuals of native species growing at equal distance in a circle (diameter = 9 cm) around each alien target plant (Figure 1). After transplanting, we randomly assigned all pots to positions on four benches of a greenhouse under natural light conditions, with a temperature between 20 and 28 °C. After five weeks, we re-randomized the positions of all pots again. To avoid leakage of nutrient solution, as well as pot-to-pot cross contamination by soil microbes, we put a plastic dish under each pot.

To test the interactive effects of nutrient availability, nutrient fluctuations and soil microbes on alien plant invasion into resident native communities, we assigned the 40 pots of each alien target species to two levels of soil-microbe (living *vs* sterilized) treatments fully crossed with two levels of nutrient-availability (low *vs* high) and two levels of nutrient-fluctuation (constant *vs* pulsed) treatments. In other words, per alien species, we had five pots (i.e., replicates), each with a different native community, in each of the eight treatment combinations. To create different soil-microbe treatments, we had filled each pot with a substrate mixture of 37.5% (v/v) sand, 37.5% (v/v) vermiculite and 25% living or sterilized field soil. The living field soil served as inoculum to provide a soil microbial community, and was collected from the surface soil (0-20 cm depth) of a grassland site in Jinlin province (44°0’4”N, 125°24’12”E), China, on 21 August 2020. We sieved the field soil through a metal grid with a mesh size of 0.5 cm, and thereafter stored half of the soil at 4 °C until pot-filling. The other half of the soil we sterilized with a dose of 25 kGy of 60CO_γ_ irradiation for four days at the Harbin Guangya Radiation New Technology Co., Ltd (Harbin, China). On 3 September 2020 (i.e., four days after transplanting), we started to apply the different nutrient treatments at weekly intervals for a total of ten weeks (Fig. 1). We applied the four nutrient treatments (2 nutrient availabilities × 2 nutrient fluctuations) using a 400%-strength Hoagland solution (for recipe details see Ref. ^8^). During the 10 weeks of the experiment, we added a total of 50 ml and 400 ml of the Hoagland solution per pot in the low and high nutrient-availability treatment, respectively. Within each nutrient-availability treatment, there were two different nutrient-supply patterns, the constant nutrient supply and the pulsed nutrient supply (Fig. 1). For the constant treatment, we supplied each pot every week with 5 ml and 40 ml of the nutrient solution for the low and high nutrient-supply treatments, respectively. For the pulsed treatment at low-nutrient availability, we first supplied each pot with 2 ml of the nutrient solution per week for three weeks, then with 9.5 ml for four weeks, and again with 2 ml for the last three weeks (Fig. 1). For the pulsed treatment at high-nutrient availability, we first supplied each pot with 10 ml of the nutrient solution per week for three weeks, then with 85 ml for four weeks, and again with 10 ml for the last three weeks (Fig. 1). To avoid that the plants would receive different amounts of water, we added extra water to the nutrient solution in each specific nutrient-supply treatment to ensure that each pot received a total of 85 ml water per nutrient application. To avoid water limitation, we checked the pots every day and watered all plants when needed by filling the dishes under the pots.

### Plant harvest and measurements

On 13 November 2020 (i.e., c. 11 weeks after transplanting), we started to harvest the aboveground biomass of all pots. We first harvested the alien target species and then harvested the native community of each pot. As one alien target plants had been planted incorrectly and 10 pots had accidentally been treated with the wrong nutrient solution, we only harvested 309 of the 320 pots. All aboveground biomass samples were dried at 65 °C for 72 hours and then weighed. Based on the aboveground biomass, we calculated the total aboveground biomass production per pot (i.e., alien target species + native community) and the biomass proportion of the alien target species in each pot (i.e., alien target species / [alien target species + native community]).

### Soil sampling, DNA extraction, amplicon sequencing and bioinformatics

To gain insights into how effects of nutrient availability and fluctuations therein on plants may be mediated by soil microbes, we also tested how the nutrient treatments influenced the soil-microbial communities of the pots with living soil inoculum. On 14 November 2020 (i.e., one day after plant harvest), we first homogenized the soil in each of these pots, and then collected from each pot a random soil sample of 12 ml, which we put into sterile centrifuge tubes (15 ml). The 154 soil samples were immediately stored at −80 □ until DNA extraction.

The DNA extraction, PCR amplifications and amplicon sequencing were performed by Novogene Co., Ltd (Beijing, China). Total genomic DNA was extracted from 0.5 g of each soil sample using the Magnetic Soil And Stool DNA Kit (Tiangen Biotech [Beijing] Co., Ltd) following the cetyltrimethylammonium bromide protocol ^19^. Then, the universal primers 341F (5’-CCTAYGGGRBGCASCAG-3’) and 806R (5’-GGACTACNNGGGTATCTAAT-3’) were used to amplify the V3-V4 region of the bacterial 16S rRNA gene. The specific fungal primers ITS1-1F-F (5’-GGAAGTAAAAGTCGTAACAAGG-3’) and ITS1-1F-R (5’-GCTGCGTTCTTCATCGATGC-3’) were used to amplify the ribosomal internal transcribed spacer of the ITS rDNA gene. All PCR reactions were carried out with 15 μL of Phusion^®^ High-Fidelity PCR Master Mix (New England Biolabs) under the following conditions: initial denaturation at 98 □ for 1 min, followed by 30 cycles of denaturation at 98 □ for 10 s, annealing at 50□ for 30 s and elongation at 72 □ for 30 s, and a final extension step of 5 min at 72 °C. PCR products were mixed in equidensity ratios. Then, mixtures of PCR products were purified with the Qiagen Gel Extraction Kit (Qiagen, Germany). Sequencing libraries were generated using the TruSeq^®^ DNA PCR-Free Sample Preparation Kit (Illumina, USA), and its quality was assessed on a Qubit@ 2.0 Fluorometer (Thermo Scientific) and Agilent Bioanalyzer 2100 system. At last, the library was sequenced on an Illumina NovaSeq platform (Illumina, USA) and 250 bp paired-end reads were generated. As fungal DNA extraction failed for one sample, we finally had 153 samples of the ITS rDNA gene, and 154 samples of the 16S rRNA gene.

The raw paired-end reads were assigned to samples, truncated by removing the barcode and primer sequence, merged using FLASH (V1.2.7, http://ccb.jhu.edu/software/FLASH/) to get splicing sequences. According to the QIIME (Quantitative Insights Into Microbial Ecology, V1.9.1, http://qiime.org/scripts/split_libraries_fastq.html) ^20^, the raw reads were quality filtered under specific filtering conditions to obtain high-quality clean reads. The clean reads were obtained after detecting and removing the chimera sequence by UCHIME algorithm ^21^. All effective reads were clustered and classified into the same OTUs (Operational Taxonomic Units) with an identity of 97% similarity using the UPARSE software (Uparse v7.0.1001; http://www.drive5.com/uparse/). The Mothur method and the SSUrRNA database ^22^ of SILVA138 (http://www.arb-silva.de) were used to annotate taxonomic information for each representative sequence of bacteria (with a threshold value of 0.8-1). The blast method ^23^ and the UNITE database (v8.2, https://unite.ut.ee/) ^24^ were used for sequence annotation analysis to obtain taxonomic information of fungal OTUs. To account for differences in sequencing depths, bacteria and fungi were rarefied to 38,678 and 47,773 reads, respectively, using the RAM package ^25^. Putative fungal functional groups (e.g., AMF and plant pathogens) with “highly probable” and “probable” confidence were identified using the FUNGuild database (v1.1) ^26^.

### Statistical analyses

To test the effects of the nutrient-availability, nutrient-fluctuation and soil-microbe treatments, and their interactions on aboveground biomass production of the alien target species, biomass production of the native communities and biomass proportion of the alien target species in each pot, we fitted Bayesian multilevel models using the function brm of the R package brms ^27^ in R 4.0.2 ^28^. To improve normality and homoscedasticity of the residuals, aboveground biomass production of the alien target species and biomass production of the native communities were natural-log-transformed, and biomass proportion of the alien target species in each pot was logit-transformed prior to analyses. In all models, we included the nutrient-availability (low *vs* high), nutrient-fluctuation (constant *vs* pulsed) and soil-microbe (living *vs* sterilized) treatments, and all their interactions as fixed factors.

To test the effects of nutrient availability, nutrient fluctuations and their interaction on the true Shannon diversity (i.e., effective number of species; ^29–32^ of soil bacteria, soil fungi and soil pathogenic fungi, as well as on the ratio between the diversity of pathogenic fungi and the diversity of AMF, we also fitted Bayesian multilevel models using the function brm of the R package brms ^27^ in R 4.0.2 ^28^. The true Shannon diversity of soil bacteria, all soil fungi and soil pathogenic fungi were calculated by the function true.diversity of the RAM package ^25^. To improve normality and homoscedasticity of the residuals, the diversity of complete soil fungi, soil pathogenic fungi and the ratio between the diversity of pathogenic fungi and AMF were natural-log-transformed prior to analyses. In all models, we included nutrient availability (low *vs* high), nutrient fluctuations (constant *vs* pulsed) and their interaction as fixed factors.

For all models mentioned above, we included identities of the plant family and the alien target species as random factors to account for non-independence of species belonging to the same family and for non-independence of replicates of the same species. In addition to account for variation among the five different native communities, we also included identity of the native community as a random factor in all models. To relax the homogeneity of variance assumption in all models, we allowed the residual standard deviation sigma to vary by the identity of the alien target species ^33^. We used the default priors set by the brms package, and ran four independent chains for all models. The total number of iterations per chain was 8000, and the number of warm-up samples was 4000. To directly test hypotheses about the main effects and interactive effects based on each coefficient’s posterior distribution, we used the sum coding, which effectively ‘centers’ the effects to the grand mean (i.e., the mean value across all data observations) ^34^. To implement this in brms, we used the functions contrasts and contr.sum of the stats package in R. We considered the fixed effects nutrient availability, nutrient fluctuations and soil microbes, and their interactions as significant when their 95% credible interval of the posterior distribution did not overlap zero, and as marginally significant when the 90% credible interval did not overlap zero.

To visualize and analyse any effects of the nutrient treatments on the composition of the microbial soil communities, we did non-metric multidimensional scaling (NMDS) based on Bray-Curtis dissimilarity distances and permutational analysis of variance (PERMANOVA), using the functions metaMDS and adonis of the vegan package in R 4.0.2 ^28^.

## Results

### Plant performance

Averaged across the nutrient-fluctuation and soil-microbe treatments, an increase in nutrient availability significantly enhanced the biomass production of alien target species (+98.2%) and the biomass production of native communities (+142.2%), but decreased the relative biomass production of target alien species in the native communities (−13.9%; Extended data Table 1; Fig. 2 and Fig. S1). Although a pulsed nutrient supply did not significantly affect the biomass production of alien target species and native communities, it tended to increase the relative biomass production of alien target species (+9.7%; 90% CIs: [0.009, 0.157] in Extended data Table 1; Fig. 2 and Fig. S1). Inoculation with living soil significantly decreased the biomass production of native communities (Fig. 2 and Fig. S2), particularly under low nutrient availability (low *vs* high: −37.6% *vs* −1.6%; Extended data Table 1; Fig. 2 and Fig. S3), and significantly increased the biomass production of alien target species (+28.8%; Extended data Table 1; Fig. 2 and Fig. S2). Consequently, the living soil microbiome significantly increased the relative biomass production of alien target species (Fig. 2 and Fig. S2), in particular under low nutrient availability (low *vs* high: +55.8% *vs* +15.1 %; Extended data Table 1; Fig. 2 and Fig. S3).

**Figure 2.**
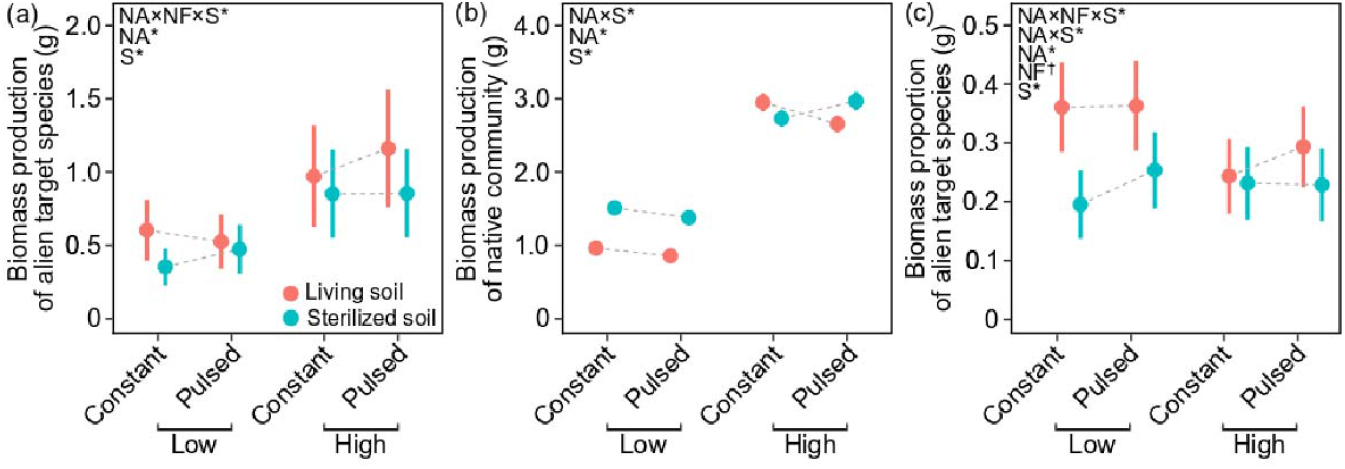
Modelled mean values (±SE) of biomass production of the alien target species (a) and the native communities (b), and biomass proportion of the alien target species (c) under each combination of two nutrient-availability (low *vs* high), two nutrient-fluctuation (constant *vs* pulsed) and two soil-microbe (living *vs* sterilized) treatments. Model terms (NA: nutrient availability, NF: nutrient fluctuation, S: soil microbiome) whose 95% credible intervals do not overlap with zero are indicated with asterisks (*), and those whose 90% credible intervals do not overlap with zero are indicated with daggers (†).

For biomass production of the alien target species, we also found a significant three-way interaction of the soil-microbe, nutrient-availability and nutrient-fluctuation treatments (Extended data Table 1). Specifically, when plants grew on substrate with sterilized soil inoculum, the pulsed nutrient supply tended to increase the biomass production of alien target species when the overall nutrient availability was low (+35.2%), whereas its effect was minimal when the overall nutrient availability was high (+0.5%; Fig. 2). However, when plants grew on substrate with living soil inoculum, the pulsed nutrient-supply effect tended to decrease the biomass production of alien target species when the overall nutrient availability was low (−13.3%), whereas the opposite was true when the overall nutrient availability was high (+19.5 %; Fig. 2). Another three-way interaction was found for the relative biomass production of alien target species (Extended data Table 1). When plants grew on substrate with sterilized soil inoculum, the pulsed nutrient supply tended to increase the relative biomass production of alien target species when the overall nutrient availability was low (+27.2%), whereas its effect was minimal when the overall nutrient availability was high (−1.2%; Fig. 2). However, when plants grew with living soil inoculum, this pattern was reversed (pulsed nutrient-supply effect under low *vs* high nutrient availability: +0.5% *vs* +19.1%; Fig. 2).

### Soil microbial community

For the soil bacterial community, we did not find any significant effects of nutrient availability, nutrient fluctuations and their interaction on the true Shannon diversity (Extended data Table 2; Fig. 3). For the complete soil fungal community, an increase in nutrient availability decreased the true Shannon diversity significantly (−14.6%; Extended data Table 2; Fig. 3 and Fig. S4). For the subset of soil pathogenic fungi, an increase in nutrient availability significantly decreased the true Shannon diversity (Fig. 3 and Fig. S4), particularly under a constant nutrient supply (constant vs pulsed: −14.9% *vs* −1.8%; Extended data Table 2; Fig. 3). In other words, a constant nutrient supply decreased the diversity of soil pathogenic fungi under high nutrient availability, but increased it under low nutrient availability. As we found no significant effects of nutrient treatments on the diversity of AMF (Fig. S4), the pulsed nutrient supply consequently tended to decrease the diversity of soil pathogenic fungi relative to AMF under low nutrient availability (−20.3%), whereas the reserve was true under high nutrient availability (+51.1%; Fig. 3). This was reflected in a marginally significant two-way interaction between nutrient availability and fluctuations (90% CIs: [0.002, 0.222] in Extended data Table 2). In line with the abovementioned findings, non-metric multidimensional scaling analysis (NMDS) and permutational multivariate analysis of variance (PERMANOVA) also showed that the nutrient treatments affected the soil microbial community composition significantly (Table S2; Extended Data Fig. 1).

**Figure 3.**
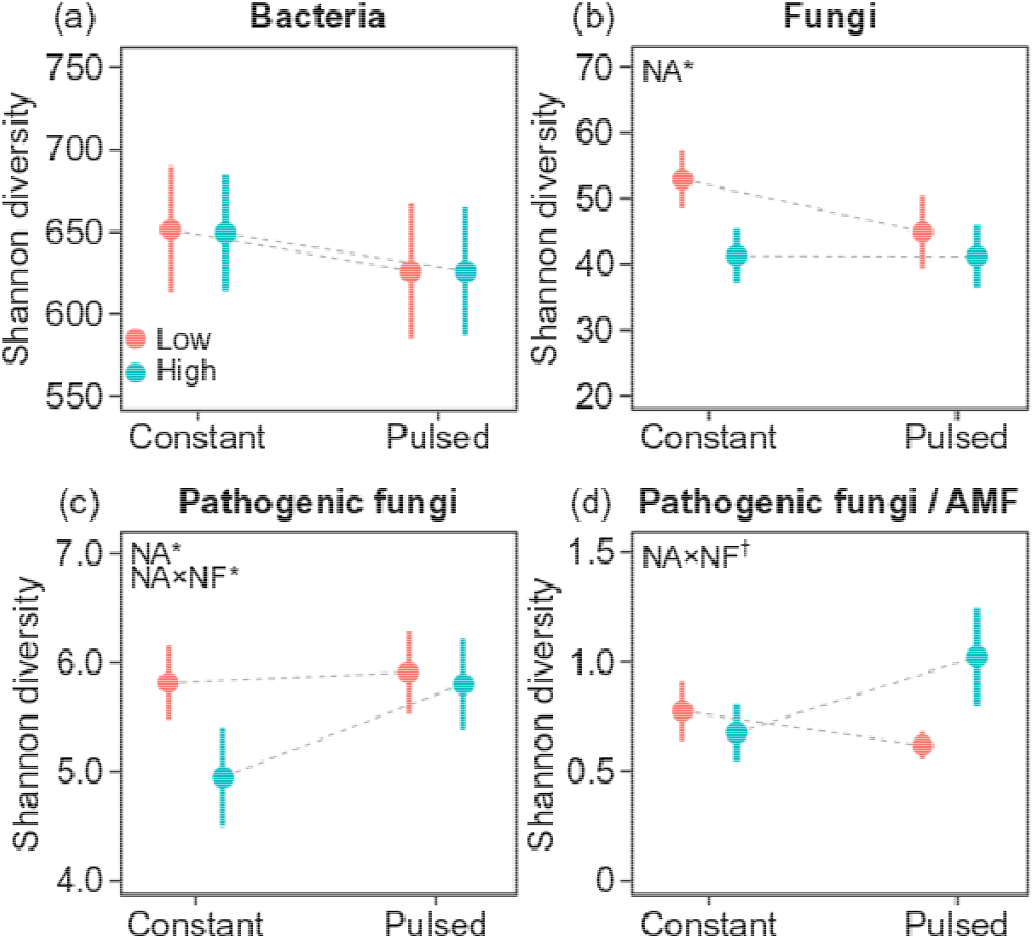
Mean values (±SE) of the true Shannon diversity for soil bacteria (a), all soil fungi (b) and the subset of pathogenic soil fungi (c), as well as the ratio of the Shannon diversity of pathogenic soil fungi and AMF (d) under each combination of two nutrient-availability (low *vs* high) and two nutrient-fluctuation (constant *vs* pulsed) treatments. Model terms (NA: nutrient availability, NF: nutrient fluctuation) whose 95% credible intervals do not overlap with zero are indicated with asterisks (*), and those whose 90% credible intervals do not overlap with zero are indicated with daggers (†).

## Discussion

Our multispecies experiment showed that the presence of a living soil microbiome increased the dominance of alien plants in the native communities. An increase in average nutrient availability also promoted the dominance of naturalized alien plants, but only when they grew on substrate with sterilized soil inoculum. Interestingly, we found that when plants grew in the absence of soil microbes, nutrient fluctuations promoted the dominance of alien plants under low average nutrient availability, but hardly affected it under high average nutrient availability. However, opposite patterns were found when plants grew in the presence of soil microbes. These different patterns may be driven by the fact that nutrient fluctuations increased the diversity of pathogenic soil fungi under high nutrient availability, but not under low nutrient availability. Our findings thus partly support the idea that environmental variability promotes plant invasion, and indicate that pathogenic soil fungi could mediate the interactive effect of nutrient availability and fluctuation therein on the dominance of alien plants in native communities.

In support of the predictions of the enemy release hypothesis, we found that the presence of soil microbes decreased the biomass production of native communities while there was a positive effect on the alien target species. As a consequence the soil microbes increased the dominance of the alien plants. This most likely reflects that the alien plants escaped from heavy pathogen loads in their native range, whereas the native communities still suffered from their soil microbial pathogens ^15,35,36^. Our results also showed that an increase in nutrient availability significantly decreased the diversity of pathogenic soil fungi. This decrease in pathogen diversity may have led to a stronger biomass increase for native communities (+142.2%) than for alien target species (+98.2%) in response to an increase in nutrient availability. As a consequence, the increase in nutrient availability tended to suppress the dominance of alien target species in the pots with living soil inoculation. However, when soil microbes were absent, our finding is in line with many theoretical ^6,37^ and empirical studies ^8,38^ showing that an increase in nutrient availability promotes alien plant invasion in native communities. These findings underline that integrating disparate invasion hypotheses will help to better understand alien invasions ^39^.

Due to higher growth rates, naturalized alien species are often more dominant than native species ^40^. Our results corroborate this, and showed that although alien plant accounted for only one-sixth of the number of individuals per pot, they accounted for about one-third of the biomass production. We also found that, averaged over all treatments, this dominance tended to increase with a pulsed nutrient supply. This finding supports the prediction of the fluctuating resource hypothesis ^6^ and is in line with results of previous empirical studies ^7,12^. However, it should be noted that in our experiment, the effect of nutrient fluctuations on alien plant invasion depended on the interaction between nutrient-availability and the presence of soil microbes.

When plants grew in substrate with sterilized soil inoculum, the dominance of alien plants was, as expected, increased by the pulsed nutrient supply under low nutrient availability. However, its effect was minimal under high nutrient availability. This most likely reflects that native species were also released from the intense belowground alien-native competition under a relative high overall nutrient availability. Consequently, the five native plants in each pot were able to take advantage of the pulsed supply pattern under high average nutrient availability, and thus their relative biomass production increased and the dominance of the alien target species was suppressed. When plants grew in substrate with living soil inoculum, however, the dominance of alien plants was only increased by the pulsed nutrient supply under high average nutrient availability. The response pattern of aliens’ dominance in response to nutrient fluctuations coincides with the changes of soil pathogen diversity, as well as the relative diversity changes between soil pathogens and AMF. A possible explanation is that the increase of soil-pathogen diversity and the relative decrease of mutualism diversity induced by the nutrient pulse under high nutrient availability suppressed the native communities but benefited the alien target species. As a consequence, alien plant dominance was promoted under these conditions. Similarly, soil pathogens most likely also mediated the effects of nutrient fluctuations on alien plant invasion when plants grew under low nutrient availability.

Our finding that soil pathogenic fungi could significantly mediate the effects of environmental variability on plant invasion has several implications. First, it suggests that the findings of previous studies that only considered direct effects of environmental variability — and nutrient fluctuations in particular— on alien plant invasion need to be further verified after accounting for indirect effects of soil microbes. Second, besides the pathogenic soil microbes, many other enemies, such as various above- and below-ground herbivores, as well as mutualisms, such as AMF and pollinators, may also indirectly steer global change driven alien plant invasion. Therefore, with the forecasted increase in environmental variability ^3^, more studies are needed to test the indirect effects of environmental variability on alien plant invasion via biotic factors.

## Conclusions

The fluctuating resource hypothesis suggests that resource fluctuations could promote alien plant invasion in native communities. Our study suggests that this is particularly the case when plants grow in the absence of soil microbes and under relatively low nutrient conditions. This finding highlights that besides the direct influence, environmental variability may also indirectly affect alien plant invasion via biotic factors such as soil microbes. Moreover, our results emphasize that to better understand alien plant invasion, integration of different invasion hypotheses is needed.

## Acknowledgements

We thank Huifei Jin, Lichao Wang, Yanjun Li, Mingxin Pan and Lichao Wang for help with the set-up of the experiment, plant harvest and biomass weighing. YL acknowledges funding from the Chinese Academy of Sciences (Y9B7041001).

## Author contributions

YL conceived the idea and designed the experiment. XZ performed the experiment. XZ, CC and YL analyzed the data. XZ and YL wrote the draft of the manuscript, with major inputs from MvK, and further inputs from CC.

## Data accessibility

Should the manuscript be accepted, the data supporting the results will be archived in Dryad and the data DOI will be included at the end of the article.

## Extended data Tables

**Extended data Table 1.**
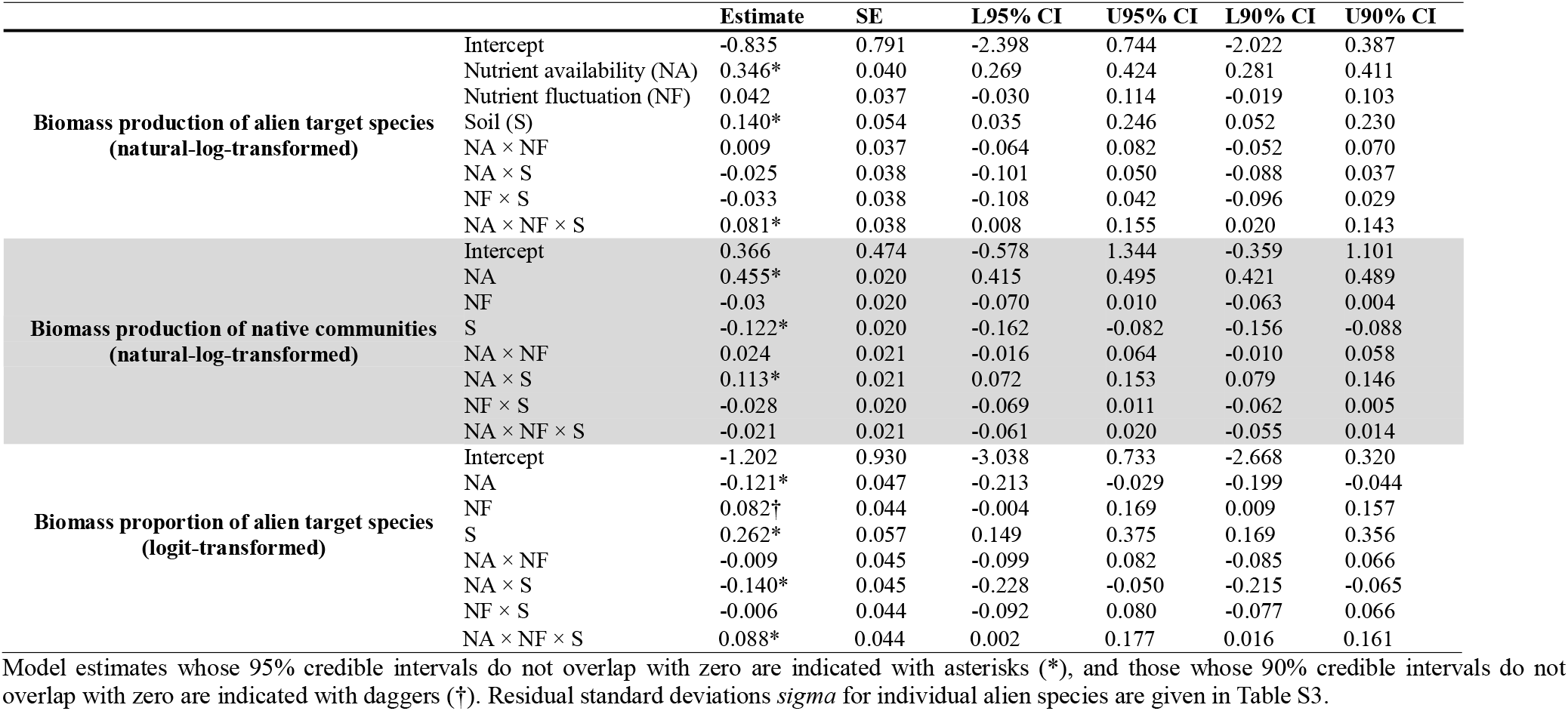
Output of the Bayesian multilevel models testing effects of the nutrient-availability (low *vs* high), nutrient-fluctuation (constant *vs* pulsed) and soil-microbe (living *vs* sterilized) treatments, and their interactions on aboveground biomass production of the alien target species, aboveground biomass production of the native communities and aboveground biomass proportion of the alien target species in each pot. Shown are the model estimates and standard errors (SE), as well as the lower (L) and upper (U) values of the 95% and 90% credible intervals (CI).

**Extended data Table 2.** Output of the Bayesian multilevel models testing effects of the nutrient-availability (low *vs* high) and nutrient-fluctuation (constant *vs* pulsed) treatments, and their interactions on the true Shannon diversity of soil bacteria, all soil fungi and the subset of pathogenic soil fungi, as well as the ratio of the Shannon diversity of pathogenic soil fungi and AMF in each pot. Shown are the model estimates and standard errors (SE) as well as the lower (L) and upper (U) values of the 95% and 90% credible intervals (CI).

|  |  | Estimate | SE | L95% CI | U95% CI | L90% CI | U90% CI |
| --- | --- | --- | --- | --- | --- | --- | --- |
| **Shannon diversity of soil bacteria** | Intercept | 639.536* | 41.416 | 557.021 | 725.848 | 575.918 | 706.156 |
|  | NA | -2.444 | 18.923 | -39.929 | 34.828 | -33.037 | 28.477 |
|  | NF | -4.817 | 18.811 | -42.377 | 31.883 | -36.117 | 25.728 |
|  | NA×NF | 0.035 | 18.425 | -36.669 | 35.901 | -29.945 | 30.258 |
| **Shannon diversity of soil fungi**<br>**(square-root-transformed)** | Intercept | 3.614* | 0.145 | 3.317 | 3.896 | 3.388 | 3.828 |
|  | NA | -0.115* | 0.057 | -0.228 | -0.002 | -0.210 | -0.021 |
|  | NF | -0.032 | 0.060 | -0.152 | 0.085 | -0.132 | 0.066 |
|  | NA×NF | 0.063 | 0.057 | -0.048 | 0.174 | -0.030 | 0.157 |
| **Shannon diversity of soil pathogenic fungi**<br>**(natural-log-transformed)** | Intercept | 1.617* | 0.125 | 1.376 | 1.868 | 1.424 | 1.814 |
|  | NA | -0.076* | 0.035 | -0.146 | -0.007 | -0.134 | -0.018 |
|  | NF | 0.053 | 0.035 | -0.015 | 0.122 | -0.004 | 0.110 |
|  | NA×NF | 0.075* | 0.036 | 0.003 | 0.145 | 0.015 | 0.134 |
| **Ratio of Shannon diversity**<br>**between soil pathogenic fungi and AMF**<br>**(natural-log-transformed)** | Intercept | -0.643* | 0.156 | -0.964 | -0.333 | -0.897 | -0.395 |
|  | NA | -0.057 | 0.069 | -0.195 | 0.077 | -0.169 | 0.057 |
|  | NF | 0.087 | 0.068 | -0.047 | 0.219 | -0.025 | 0.198 |
|  | NA×NF | 0.112† | 0.068 | -0.020 | 0.246 | 0.002 | 0.222 |

## Extended data Figures

**Extended data Figure 2.**
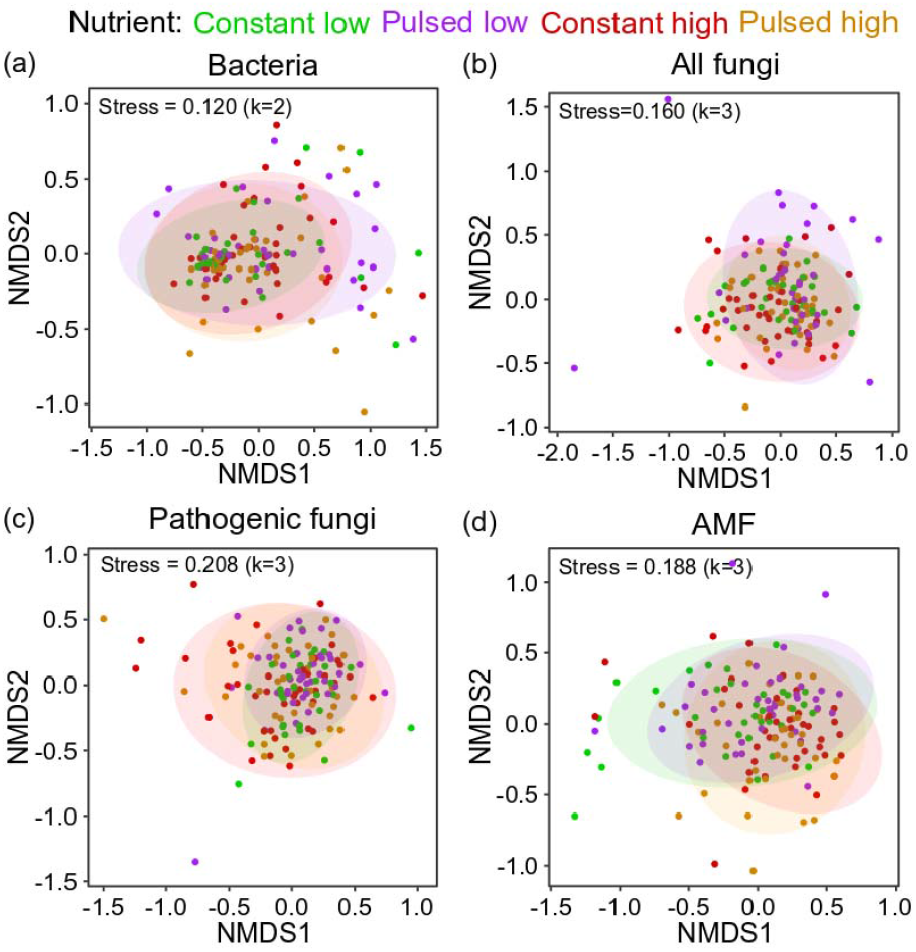
Effects of nutrient treatments on soil-community composition of bacteria (a), all fungi (b), pathogenic fungi (c) and AMF (d). Nonmetric multidimensional scaling (NMDS) was used to visualize differences in the soil microbial communities of the nutrient treatments. Data points represent soil samples. The different colors of the points indicate different nutrient treatments. Ellipses represent means ± 1 SDs for soils under constant low (green), pulsed low (purple), constant high (red) or pulsed high nutrient supply (orange).

## Supporting information

**Table S1.** Details of the study species used in the experiment.

| Species | Family | Status | Community group | Sowing date | Life history | Propagule source |
| --- | --- | --- | --- | --- | --- | --- |
| *Bidens frondosa* L. | Compositae | Alien | / | 9.08.2020 | Annual | wild population |
| *Bidens pilosa* L. | Compositae | Alien | / | 09.08.2020 | Annual | wild population |
| *Erigeron annuus* (L.) Pers. | Compositae | Alien | / | 26.06.2020 | Annual/Biennial | wild population |
| *Erigeron canadensis* L. | Compositae | Alien | / | 26.06.2020 | Annual | wild population |
| *Lolium perenne* L. | Poaceae | Alien | / | 14.08.2020 | Perennials | commercial |
| *Paspalum notatum* Flügge | Poaceae | Alien | / | 06.08.2020 | Perennials | wild population |
| *Phytolacca americana* L. | Phytolaccaceae | Alien | / | 26.06.2020 | Perennials | commercial |
| *Solidago canadensis* L. | Compositae | Alien | / | 03.08.2020 | Perennials | wild population |
| *Cynanchum chinense* R.Br. | Apocynaceae | Native | NC1 | 14.08.2020 | Annual | wild population |
| *Digitaria sanguinalis* (L.) Scop | Poaceae | Native | NC1 | 07.08.2020 | Annual | wild population |
| *Potentilla tanacetifolia* Willd. ex Schtdl. | Rosaceae | Native | NC1 | 07.08.2020 | Perennials | wild population |
| *Dontostemon micranthus* C.A.Mey. | Brassicaceae | Native | NC1 | 07.08.2020 | Annual | wild population |
| *Cleistogenes squarrosa* (Trin. ex Ledeb.) Keng | Poaceae | Native | NC1 | 11.08.2020 | Perennials | wild population |
| *Echinochloa crus-galli* (L.) P.Beauv. | Poaceae | Native | NC2 | 07.08.2020 | Annual | wild population |
| *Artemisia gmelinii* Weber ex Stechm. | Compositae | Native | NC2 | 07.08.2020 | Perennials | wild population |
| *Crepidiastrum sonchifolium* (Maxim.) Pak & Kawano | Compositae | Native | NC2 | 07.08.2020 | Annual/Biennial | wild population |
| *Eragrostis pilosa* (L.) P.Beauv. | Poaceae | Native | NC2 | 14.08.2020 | Annual | wild population |
| *Gypsophila licentiana* Hand. Mazz. | Caryophyllaceae | Native | NC2 | 07.08.2020 | Perennials | wild population |
| *Geum aleppicum* Jacq. | Rosaceae | Native | NC3 | 07.08.2020 | Perennials | wild population |
| *Lappula myosotis* V. Wolf | Boraginaceae | Native | NC3 | 07.08.2020 | Annual/ Biennial | wild population |
| *Chloris virgata* Sw. | Poaceae | Native | NC3 | 14.08.2020 | Annual | wild population |
| *Euphorbia humifusa* Willd. | Euphorbiaceae | Native | NC3 | 07.08.2020 | Annual | wild population |
| *Bromus inermis* Leyss | Poaceae | Native | NC3 | 07.08.2020 | Perennials | wild population |
| *Malva verticillata* L. | Malvaceae | Native | NC4 | 07.08.2020 | Biennial | wild population |
| *Achillea asiatica* Serg. | Compositae | Native | NC4 | 07.08.2020 | Perennials | wild population |
| *Poa annua* L. | Poaceae | Native | NC4 | 07.08.2020 | Annual | wild population |
| *Dysphania schraderiana* (Schult.) Mosyakin & Clemants | Amaranthaceae | Native | NC4 | 07.08.2020 | Annual | wild population |
| *Setaria pumila* (Poir.) Roem. & Schult. | Poaceae | Native | NC4 | 14.08.2020 | Annual | wild population |
| *Rumex crispus* L. | Polygonaceae | Native | NC5 | 07.08.2020 | Perennials | wild population |
| *Setaria viridis* (L.) P.Beauv. | Poaceae | Native | NC5 | 14.08.2020 | Annual | wild population |
| *Artemisia rubripes* Nakai | Compositae | Native | NC5 | 11.08.2020 | Perennials | wild population |
| *Eragrostis minor* Host | Poaceae | Native | NC5 | 14.08.2020 | Annual | wild population |
| *Potentilla longifolia* Willd. ex Schtdl. | Rosaceae | Native | NC5 | 07.08.2020 | Perennials | wild population |

**Table S2.** Effects of nutrient treatments on soil-community composition of bacteria, all fungi, pathogenic fungi and AMF.

|  | Bacteria |  |  |  | All fungi |  |  |  | Pathogenic fungi |  |  |  | AMF |  |  |  |
| --- | --- | --- | --- | --- | --- | --- | --- | --- | --- | --- | --- | --- | --- | --- | --- | --- |
|  | Df | F | R<sup>2</sup> | P | Df | F | R<sup>2</sup> | P | Df | F | R<sup>2</sup> | P | Df | F | R<sup>2</sup> | P |
| Nutrient availability (NA) | 1 | 3.213 | 0.021 | **0.010\*** | 1 | 2.944 | 0.019 | **0.001\*** | 1 | 3.989 | 0.026 | **0.002\*** | 1 | 5.514 | 0.035 | **0.001\*** |
| Nutrient fluctuation (NF) | 1 | 1.162 | 0.008 | 0.221 | 1 | 0.855 | 0.006 | 0.669 | 1 | 0.750 | 0.005 | 0.597 | 1 | 0.598 | 0.004 | 0.789 |
| NA × NF | 1 | 1.393 | 0.009 | 0.147 | 1 | 0.967 | 0.006 | 0.405 | 1 | 1.530 | 0.010 | 0.091† | 1 | 1.456 | 0.009 | 0.138 |
| Residuals | 150 |  | 0.963 |  | 149 |  | 0.969 |  | 149 |  | 0.960 |  | 149 |  | 0.952 |  |
| Total | 153 |  | 1.000 |  | 152 |  | 1.000 |  | 152 |  | 1.000 |  | 152 |  | 1.000 |  |

Significant effects ( $P < 0.05$ ) are indicated with asterisks (\*), and marginally significant effects ( $0.05 \leq P < 0.1$ ) are indicated daggers (†).

**Table S3.** Output of the Bayesian multilevel models’ estimates of the residual standard-deviation sigmas for individual alien species. Shown are the model estimates and standard errors as well as the lower (L) and upper (U) values of the 95% and 90% credible intervals (CI).

|  | Species | Estimate | SE | L95% CI | U95% CI | L90% CI | U90% CI |
| --- | --- | --- | --- | --- | --- | --- | --- |
| **Biomass production of alien target species** | *Bidens frondosa* L. | -0.635* | 0.128 | -0.877 | -0.376 | -0.840 | -0.421 |
|  | *Bidens pilosa* L. | -0.522* | 0.124 | -0.750 | -0.267 | -0.715 | -0.310 |
|  | *Erigeron annuus* (L.) Pers. | -0.266† | 0.134 | -0.523 | 0.006 | -0.481 | -0.042 |
|  | *Erigeron canadensis* L. | -0.186 | 0.125 | -0.417 | 0.070 | -0.383 | 0.027 |
|  | *Lolium perenne* L. | -0.492* | 0.141 | -0.760 | -0.207 | -0.717 | -0.252 |
|  | *Paspalum notatum* Flügge | -0.713* | 0.136 | -0.970 | -0.436 | -0.931 | -0.484 |
|  | *Phytolacca americana* L. | 0.005 | 0.124 | -0.230 | 0.258 | -0.191 | 0.215 |
|  | *Solidago canadensis* L. | -0.253† | 0.125 | -0.487 | 0.002 | -0.450 | -0.041 |
| **Biomass production of native communities** | *Bidens frondosa* L. | -1.095* | 0.118 | -1.315 | -0.849 | -1.282 | -0.896 |
|  | *Bidens pilosa* L. | -1.032* | 0.120 | -1.258 | -0.786 | -1.221 | -0.831 |
|  | *Erigeron annuus* (L.) Pers. | -0.810* | 0.120 | -1.035 | -0.561 | -0.998 | -0.606 |
|  | *Erigeron canadensis* L. | -0.934* | 0.121 | -1.158 | -0.683 | -1.126 | -0.727 |
|  | *Lolium perenne* L. | -1.164* | 0.120 | -1.386 | -0.918 | -1.356 | -0.958 |
|  | *Paspalum notatum* Flügge | -1.106* | 0.121 | -1.332 | -0.859 | -1.299 | -0.900 |
|  | *Phytolacca americana* L. | -0.755* | 0.120 | -0.979 | -0.507 | -0.943 | -0.552 |
|  | *Solidago canadensis* L. | -1.168* | 0.121 | -1.390 | -0.917 | -1.360 | -0.962 |
| **Biomass proportion of alien target species** | *Bidens frondosa* L. | -0.457* | 0.125 | -0.689 | -0.201 | -0.656 | -0.245 |
|  | *Bidens pilosa* L. | -0.230† | 0.122 | -0.457 | 0.017 | -0.423 | -0.023 |
|  | *Erigeron annuus* (L.) Pers. | -0.021 | 0.127 | -0.259 | 0.239 | -0.224 | 0.194 |
|  | *Erigeron canadensis* L. | -0.022 | 0.124 | -0.255 | 0.228 | -0.220 | 0.186 |
|  | *Lolium perenne* L. | -0.376* | 0.133 | -0.627 | -0.109 | -0.589 | -0.153 |
|  | *Paspalum notatum* Flügge | -0.533* | 0.131 | -0.779 | -0.265 | -0.741 | -0.312 |
|  | *Phytolacca americana* L. | 0.233* | 0.121 | 0.005 | 0.479 | 0.038 | 0.440 |
|  | *Solidago canadensis* L. | -0.203 | 0.125 | -0.437 | 0.056 | -0.401 | 0.012 |

Model estimates whose 95% credible intervals do not overlap with zero are indicated with asterisks (\*), and those whose 90% credible intervals do not overlap with zero are indicated with daggers (†).

**Table S4.** Output of the Bayesian multilevel models’ estimates of the residual standard-deviation sigmas for each community. Shown are the model estimates and standard errors as well as the lower (L) and upper (U) values of the 95% and 90% credible intervals (CI).

|  | Community group | Estimate | SE | L95% CI | U95% CI | L90% CI | U90% CI |
| --- | --- | --- | --- | --- | --- | --- | --- |
| **Shannon diversity of soil bacteria** | NC1 | 5.468* | 0.140 | 5.210 | 5.753 | 5.246 | 5.706 |
|  | NC2 | 5.518* | 0.134 | 5.271 | 5.801 | 5.306 | 5.750 |
|  | NC3 | 5.332* | 0.147 | 5.056 | 5.633 | 5.099 | 5.583 |
|  | NC4 | 5.310* | 0.141 | 5.045 | 5.604 | 5.086 | 5.553 |
|  | NC5 | 5.484* | 0.135 | 5.232 | 5.759 | 5.270 | 5.711 |
| **Shannon diversity of soil fungi** | NC1 | -0.297* | 0.132 | -0.537 | -0.020 | -0.504 | -0.071 |
|  | NC2 | -0.074 | 0.133 | -0.324 | 0.202 | -0.283 | 0.152 |
|  | NC3 | -0.572* | 0.142 | -0.837 | -0.276 | -0.794 | -0.329 |
|  | NC4 | -0.278† | 0.137 | -0.533 | 0.000 | -0.493 | -0.046 |
|  | NC5 | -0.477* | 0.142 | -0.743 | -0.183 | -0.701 | -0.235 |
| **Shannon diversity of soil pathogenic fungi** | NC1 | -0.887* | 0.137 | -1.139 | -0.601 | -1.101 | -0.653 |
|  | NC2 | -0.843* | 0.137 | -1.096 | -0.561 | -1.059 | -0.611 |
|  | NC3 | -0.941* | 0.146 | -1.213 | -0.642 | -1.175 | -0.694 |
|  | NC4 | -0.906* | 0.147 | -1.178 | -0.610 | -1.137 | -0.659 |
|  | NC5 | -0.584* | 0.140 | -0.848 | -0.292 | -0.803 | -0.346 |
| **Ratio of Shannon diversity of soil pathogenic fungi and AMF** | NC1 | -0.184 | 0.137 | -0.435 | 0.095 | -0.400 | 0.048 |
|  | NC2 | -0.091 | 0.132 | -0.339 | 0.178 | -0.301 | 0.134 |
|  | NC3 | -0.234 | 0.144 | -0.501 | 0.063 | -0.461 | 0.009 |
|  | NC4 | -0.253† | 0.146 | -0.527 | 0.046 | -0.485 | -0.008 |
|  | NC5 | -0.17 | 0.138 | -0.426 | 0.115 | -0.390 | 0.065 |

Model estimates whose 95% credible intervals do not overlap with zero are indicated with asterisks (\*), and those whose 90% credible intervals do not overlap with zero are indicated with daggers (†).

**Figure S1.**
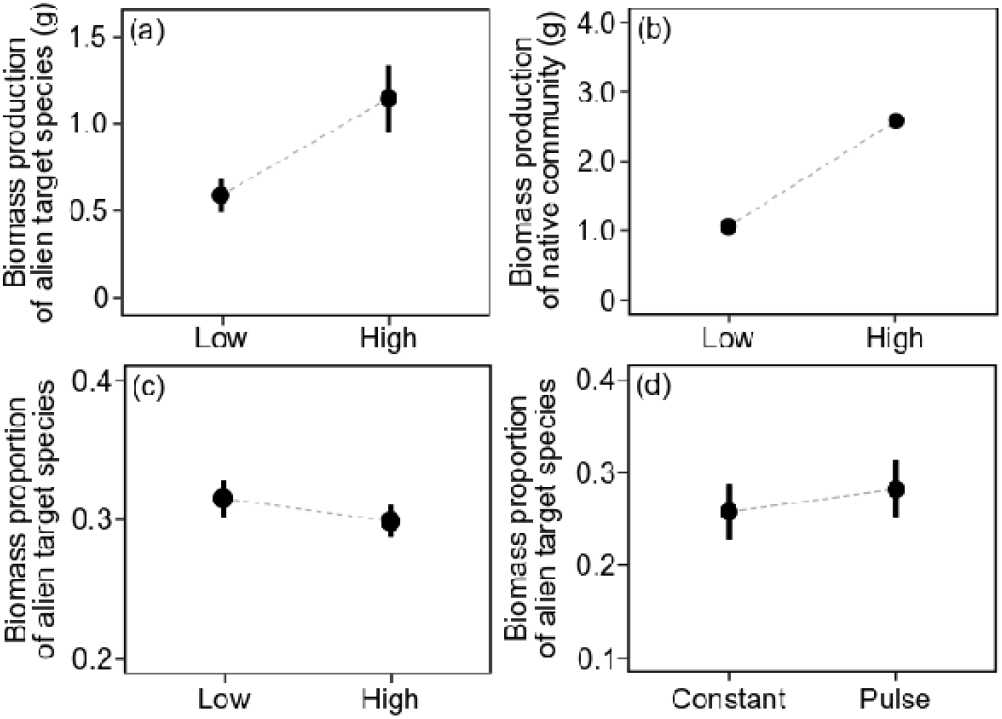
Modelled mean values (±SE) of biomass production of the alien target species (a) and the native communities (b), and biomass proportion of the alien target species (c) under low and high nutrient availability, as well as biomass proportion of the alien target species under constant and pulsed nutrient supply (d).

**Figure S2.**
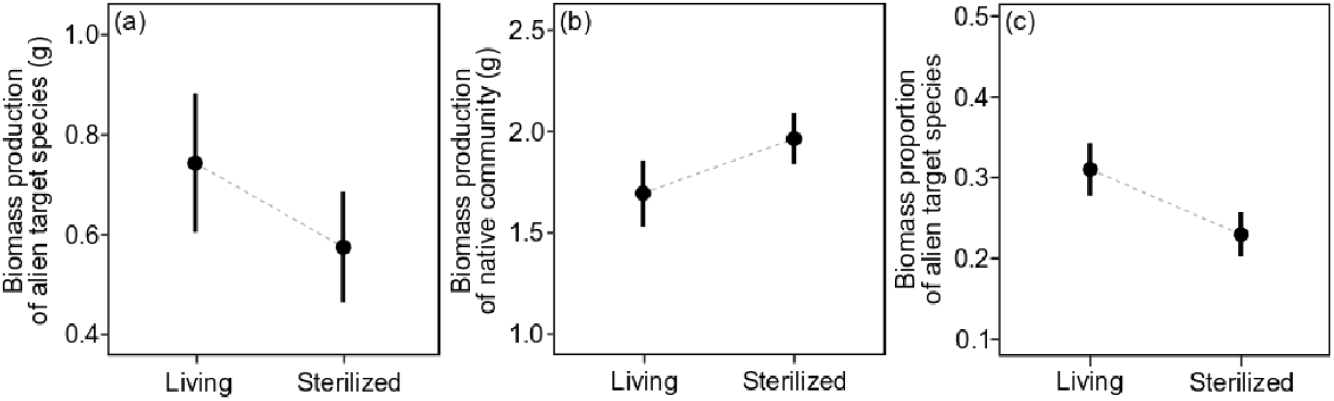
Modelled mean values (±SE) of biomass production of the alien target species (a) and the native communities (b), and biomass proportion of the alien target species (c) growing in substrate inoculated with living and sterilized field soil.

**Figure S3.**
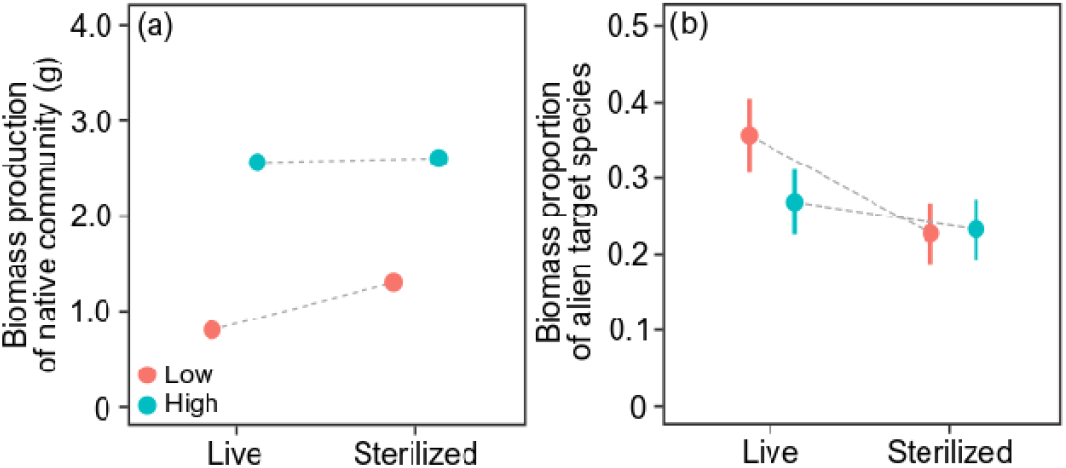
Modelled mean values (±SE) of biomass production of native communities (a) and biomass proportion of alien target species (b) under each combination of the nutrient-availability (low *vs* high) and soil-microbe (living *vs* sterilized) treatments.

**Figure S4.**
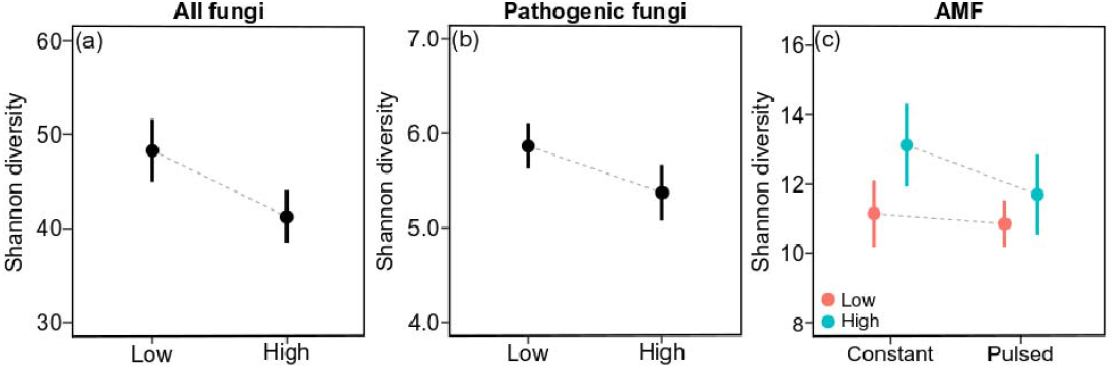
Mean values of the true Shannon diversity (±SE) of all soil fungi (a) and the subset of pathogenic soil fungi (b) under low and high nutrient availability.

